# A proteogenomics workflow to uncover the world of small proteins in *Staphylococcus aureus*

**DOI:** 10.1101/2020.05.25.114132

**Authors:** Stephan Fuchs, Martin Kucklick, Erik Lehmann, Alexander Beckmann, Maya Wilkens, Baban Kolte, Ayten Mustafayeva, Tobias Ludwig, Maurice Diwo, Josef Wissing, Lothar Jänsch, Christian H. Ahrens, Zoya Ignatova, Susanne Engelmann

## Abstract

Small proteins play diverse and essential roles in bacterial physiology and virulence. Despite their importance, automated genome annotation algorithms still cannot accurately annotate all respective small open reading frames (sORFs), as they usually provide insufficient sequence information for domain and homology searches, tend to be species specific and only a few experimentally validated examples are covered in standard proteomics studies. The accuracy and reliability of genome annotations, particularly for sORFs, can be significantly improved by integrating protein evidence from experimental approaches that enrich for small proteins. Here we present a highly optimized and flexible workflow for bacterial proteogenomics, which covers all steps from (i) creation of protein databases, (ii) database searches, (iii) peptide-to-genome mapping to (iv) result interpretation and whose automated execution is supported by two open source tools (*SALT* & *Pepper*). We used the workflow to identify high quality peptide spectrum matches (PSMs) for both annotated and unannotated small proteins (≤ 100 aa; SP100) in *Staphylococcus aureus* Newman. Proteins isolated from cells at the exponential and stationary growth phase were digested with different endopeptidases (trypsin, Lys-C, AspN), the resulting peptides fractionated by gel-based and gel-free methods and measured with highly sensitive mass spectrometers. PSMs or sORF predictions from sORFfinder were stringently filtered allowing us to detect 185 soluble SP100, 69 of which were missing in the used genome annotation. Most interestingly, almost half of the identified SP100 were basic, suggesting a role in binding to more acidic molecules such as nucleic acids or phospholipids. In addition, phage-related functions were proposed for 30 SP100, based on the localization of their coding sequences in the genome.

## 1. Introduction

*Staphylococcus aureus* is a Gram-positive human pathogen with strong clinical significance that causes mainly nosocomial infections in immunocompromised patients which are frequently associated with difficult to treat multidrug-resistant *S. aureus* phenotypes (1). With 547 complete genome sequences publically available from the National Center of Biotechnology Information (NCBI’s) Reference Sequence Database (RefSeq; status 2020-05-17), *S. aureus* is among the most sequenced bacteria. The number of thus far annotated open reading frames ranges from 2600 to 2900 per genome sequence. The entire pan-genome of *S. aureus* has not yet been described, due to the fact that the genomic diversity of *S. aureus* is very high (2,3). However, a preliminary *S. aureus* pan-genome based on the comparison of 64 *S. aureus* genome sequences is composed of 7,411 genes, of which about 20% are conserved constituting the core-genome (3). The highest variability has been found among genes coding for extracellular and surface-associated proteins (4) which is of particular importance as these proteins are essentially involved in direct interactions with the host environment during infection. The protein inventory of several *S. aureus* strains has been described using highly sensitive mass spectrometry (MS) techniques combined with liquid chromatography (LC) (5–7). For *S. aureus* strain COL, more than 1700 proteins (about 60 % of the theoretical proteome) have been identified, quantified and assigned to various subcellular localizations (5,7,8), which can help to predict functions for colocalized proteins (9). However, one group of proteins was highly underrepresented in the *S. aureus* proteome: very small proteins not longer than 100 amino acids (aa) (=SP100). Altogether, Becher and colleagues (5) detected 82 annotated SP100 of which only four proteins possess 50 aa or less (=SP50). The experimental detection of SP100 by shotgun proteomics is difficult and also affected by the fact that their encoding short open reading frames (sORFs) are frequently filtered out by conventional genome annotation algorithms. There are several reasons hindering automated annotation and prediction of sORFs, such as insufficient sequence information for domain and homology searches, a limited number of experimentally validated templates, and their tendency to species specificity (10–12). Hence, differentiation between sORFs with low and high coding potential is challenging and the number of false positives among predicted sORFs is extremely high (13). Given these facts, arbitrary cut-offs for a minimal ORF length of 50 or 100 codons were routinely used in genome annotations. In addition, the low molecular weight of these proteins complicates experimental isolation and also reduces the number of MS-compatible peptides.

Over the last years, various attempts have been made that address one or both of these issues. This includes experimental approaches such as ribosomal profiling and proteogenomics to identify these group of proteins as well as bioinformatic approaches for a more reliable prediction and comprehensive annotation of sORFs (14–26). For instance, different computational approaches have been developed for sORF prediction which have in common that the coding potential of a putative sORF is scored based on at least one or more features such as nucleotide composition, synonymous and non-synonymous substitution rates, phylogenetic conservation or protein domain detection (13,16).

Despite these major challenges, there is no doubt that small proteins play a pivotal role in essential cellular processes, hence it is extremely important to improve our ability to uncover this pool of hidden proteins (20,27–30). First functional characterizations prove their involvement in various cellular processes such as protein folding, regulation of gene expression, membrane transport, protein modification and signal transduction in different bacteria (for an overview see (30)). In addition, some small proteins have an extracellular function and exhibit toxic or antimicrobial activity. Interestingly, most of the small proteins characterized so far are associated with the cell membrane and are poorly conserved at the sequence level (29). In *S. aureus,* the most prominent small proteins are phenol soluble modulins with a length of only 20 to 40 aa (31) and delta-hemolysin (26 aa) (32). Phenol soluble modulins possess multiple roles in *S. aureus* pathogenesis by inducing cell lysis of blood cells, stimulating inflammatory responses and influencing biofilm formation (for review see (33)). Delta-hemolysin interacts with membranes of various blood cells, which concentration dependently results in a membrane disturbance and even in cell lysis (for review see (34)). While both phenol soluble modulins and delta-hemolysin have been studied in great detail in recent years, data on the identification and functional characterization of other small proteins with up to 50 aa in *S. aureus* are almost completely missing.

High availability of *S. aureus* genome sequences defines it as a well-suited model bacterium for prediction and identification of small proteins and peptides. Currently, the number of annotated coding sequences with up to 303 nucleotides is highly variable in the 547 complete *S. aureus* genome sequences and their number ranges from 345 to 632. This is mainly attributed to the fact that various algorithms of genome annotation were applied. Among *S. aureus* reference strains, strain Newman plays a pivotal role. First isolated in 1952 from a human infection (35), it is one of the most frequently used *S. aureus* strains in infection models as it is characterized by a relatively stable phenotype. In addition, four prophages were identified in the genome of strain Newman, inserted at different sites in the chromosome, exceeding the regularly observed number of prophages in *S. aureus* (36–38). In a murine infection model, the loss of all four prophages significantly reduced the virulence potential of the strain (39). The existence of these prophages made *S. aureus* Newman to an excellent model studying their impact on virulence and cells physiology. The genome sequence of strain Newman, was predicted to encode at least 2,614 proteins, again, the number of annotated proteins with 100 aa or below is rather low and amounts to XX proteins (40).

To more comprehensively identify proteins with up to 100 aa, we used *S. aureus* as a model system and developed an integrative proteogenomics workflow that combines *in silico* translation of the entire genome sequence, various LC-MS/MS workflows, and a bioinformatic pipeline for peptidomics data analyses. The workflow is highly optimized and flexible and ready to use in other bacterial species.

## 2. Results

### 2.1. Estimating the number of spurious sORFs in *S. aureus* Newman using singlenucleotide permutation testing

A single nucleotide permutation test was used to verify the global confidence and significance of ORFs based on their length only. For this purpose, ORFs were detected in the genome sequence of *S. aureus* Newman (NC_009641.1; NCBI translation table 11; longest ORFs preferred). As false-positive estimate, we used the median number of ORFs detected in permuted genome sequences (n=1000), that show the same nucleotide composition but in random order and can therefore be assumed to not contain any biological information (Fig. 1A). Accordingly, the false discovery rate (FDR) of ORFs detected in the biological sequence is 113% for those with a maximum length of 63 bp (coding for 20 aa), 133% for those with a length between 66 and 153 bp (21 to 50 aa) and 94% for those with a length between156 and 303 bp (51 to 100 aa). This highlights the need for additional evidence for the reliable annotation of sORFs. In contrast, coding sequences (CDS) with a length of at least 396 bp (≥ 131 aa) did not occur by chance in the permuted sequence set. Since start and stop codons tend to be AT-rich, the FDR for sORFs increases with an increasing GC content of an organism (Fig. 1B).

**Figure 1.**
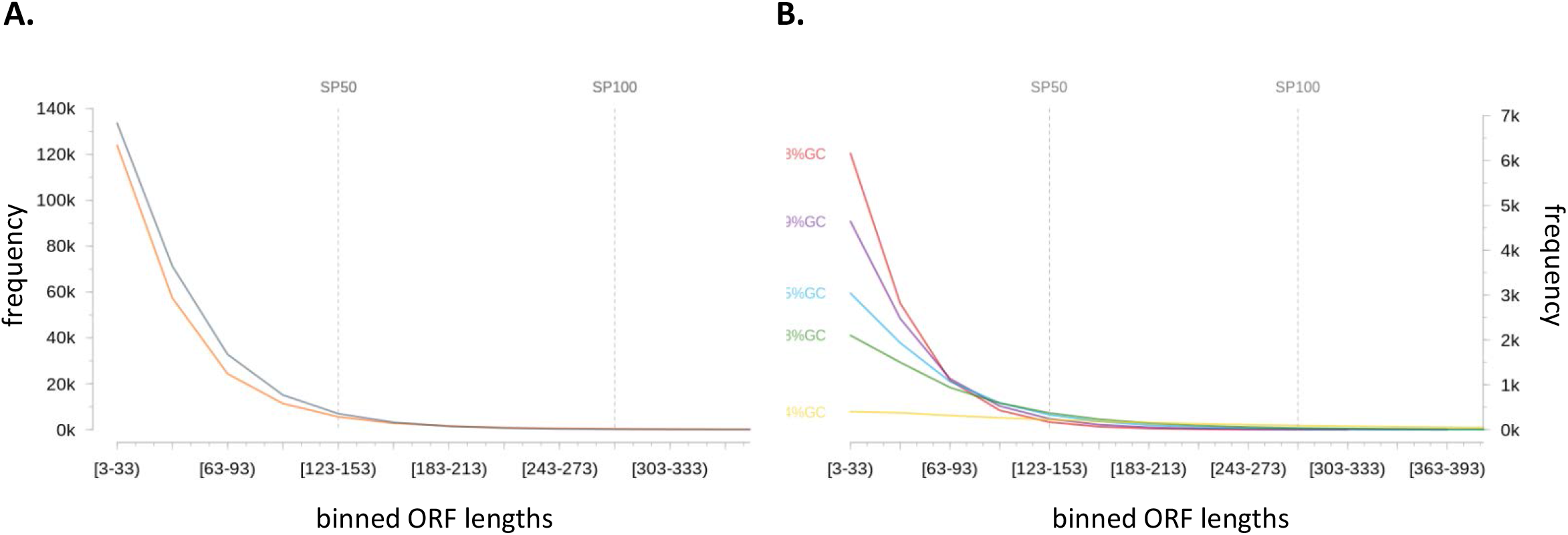
Potential sORF frequencies in biological and permuted genome sequences. **(A) Estimation of the proportion of false-positive sORFs in predictions based solely on start and stop codons:** All potential ORF sequences (NCBI translation table 11; longest ORF variants preferred) were extracted from the genome sequence of *S. aureus* Newman and 1,000 permuted sequence derivatives showing the same nucleotide composition but in random order and can therefore be assumed to no longer contain any biological information. Resulting ORFs were binned based on their length. Bin sizes are shown for the genuine reference sequence (orange) and the permuted sequences (grey; as median) up to a maximum ORF length of 303bp (= 100aa). Especially small ORFs tend to occur randomly. **(B) Impact of GC content on the number of spurious ORF:** According to (A), bin sizes are given for genome sequences and their permuted sequence derivatives (n= 1,000; as median) with varying GC content. The real sequences are from xxx.

### 2.2. Creating more comprehensive protein databases for *S. aureus* Newman using *in silico* translation and sORF prediction

To identify small proteins not covered in the RefSeq annotation, we generated protein databases using two different approaches. First, protein coding sORFs were predicted based on their coding potential using sORF finder (41). To train the classifier, all annotated CDS and sequences of stable RNAs were used as positive (coding) and negative (non-coding) templates, respectively (used annotation: NC_009641.1 as at 07/06/2013). Based on this, a high coding potential was predicted for 3,357 ORFs within intergenic regions varying between 33 and 267 bp. These ORFs were translated and combined with the annotated protein sequences resulting in a database with a total number of 6,441 sequences. In the following this database is referred to as **PR**edicted Protein **D**ata**B**ase (PRDB).

With the second approach we aimed to consider the full coding potential of *S. aureus* Newman by translating all six reading frames of the respective genome sequence and creating a separate protein entry for each sequence between two stop codons. The resulting **TR**anslation **D**ata**B**ase (TRDB) comprises 177,532 sequences with a minimum length of 9 aa. We established an automated workflow for the translation of (circular) bacterial genome sequences (Fig. 2 A). The corresponding python-based tool called *SALT* is publicly available (https://gitlab.com/s.fuchs/pepper). It allows the extraction of all potential ORFs from a circular or linear genome sequence and supports both start to stop and stop to stop codon extraction (according to NCBI translation table 11). The extracted sequences can be automatically translated and, if required, digested *in silico* into individual peptides using predefined digestion patterns of different enzymes. All DNA, protein and peptide sequences are extractable to individual FASTA files. The respective sequence header can be fully customized to meet specific requirements. Moreover, for each sequence collection, tables can be exported as tabulator delimited text files showing different physicochemical properties such as molecular weights, isoelectric points or grand average of hydropathy (GRAVY) values. To the best of our knowledge, this is the only freely available tool that offers such a variety of functions (further information see https://gitlab.com/s.fuchs/pepper).

**Figure 2.**
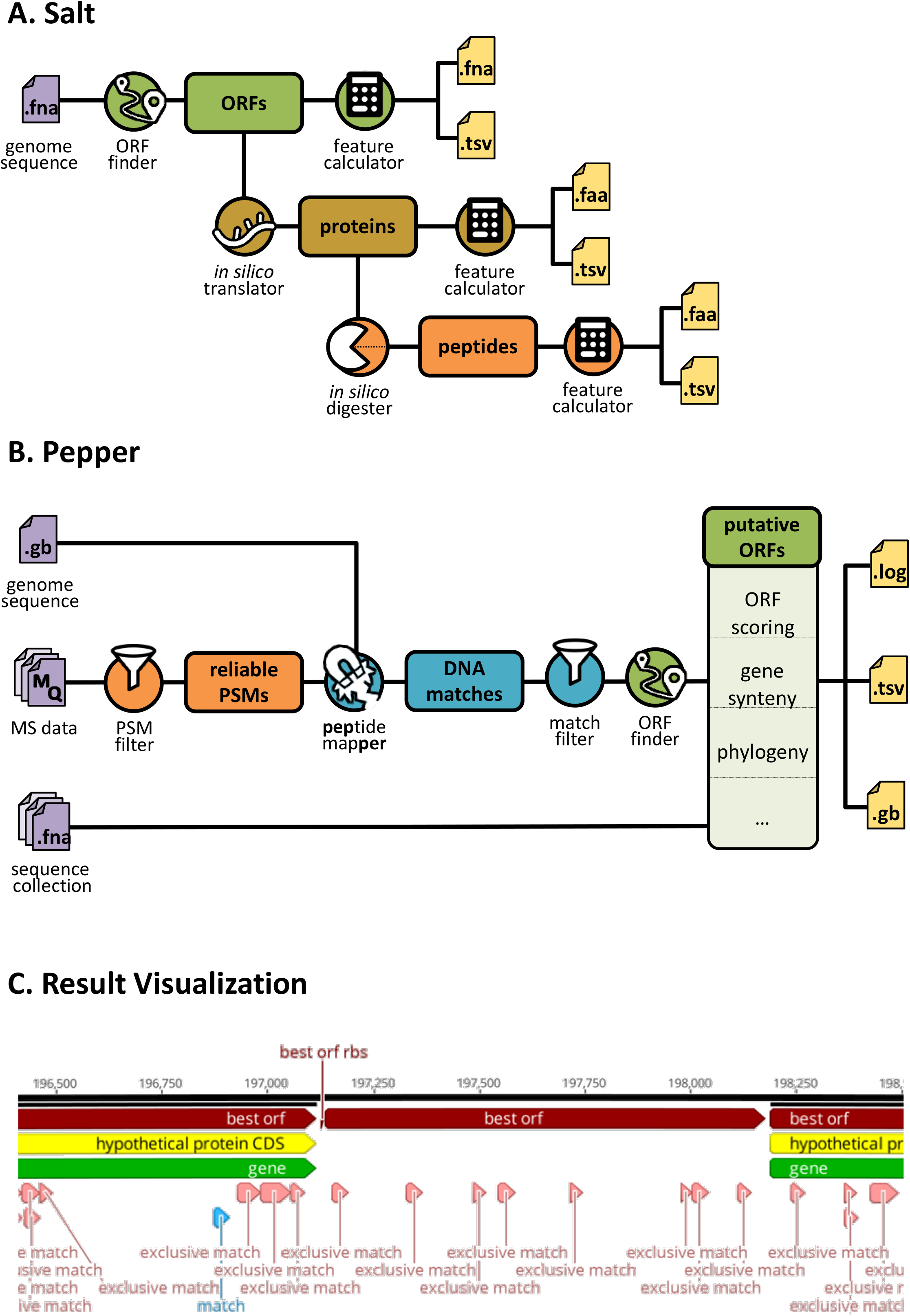
Bacterial proteogenomics workflow provided by Salt and Pepper. **(A) Creation of protein and peptide databases using Salt:** Based on a FASTA file as input, Salt extracts all potential ORFs from a given (circular) genome sequence using different methods (stop to stop codon, start to stop codon). The resulting ORF sequences are then translated *in silico* into protein sequences that can be further digested using different *in silico* proteases (Tryspin, Chymotrypsin, Asp-N, Lys-C and Proteinase K). For each level (ORFs, proteins, peptides) individual FASTA files and tables (tab-separated values, TSV) are created, listing various sequence-derived properties such as molecular weight or isoelectric points. **(B) Proteogenomics analyses using Pepper:** Peptide to spectrum matches (PSMs) obtained from different samples are directly extracted from Maxquant evidence files (MQ). Different spectrum quality and replication criteria can be defined to restrict the analysis to highly-reliable PSMs only. Respective coding sites are determined in a given (circular) genome sequence provided as FASTA file. The resulting DNA matches, that can be filtered e.g. by exclusivity, are used to predict the putative open reading frames. Additional information such as potential ribosomal binding sites, gene synteny based on the reference genome annotation, and conservation in given sequence collections (provided as FASTA files) are collected. Different files are created to archive all analysis parameters (log file), results on peptide, DNA match, and ORF level (TSV), and an updated reference genome annotation integrating the identified DNA matches and ORFs (Genbank file; GB). **(C) Results visualization:** GB files created by Pepper can be used for results visualization using third-party software (here: Geneious Prime, Biomatters Ltd.). The genome sequence (black line) with coordinates is shown on top. Existing annotations are highlighted in yellow and green. ORFs with the highest coding potential regarding Pepper and potential ribosomal binding sites (rbs) are highlighted in red. DNA matches are show in light-red, if the respective peptide is not encoded elsewhere in the genome (exclusive match), or light-blue, if multiple coding sites exist for the respective peptide.

### 2.3. Creating a fully automated yet flexible workflow for bacterial proteogenomics

To deduce putative ORFs from a list of identified peptides, we created a rule-based expert system called *Pepper* which is fully automated and optimized for bacterial proteogenomics (Fig. 2B). In brief, sequences and quality measures of peptide spectrum matches (PSMs) are extracted from evidence files provided by the Maxquant software (Max Planck Institute of Biochemistry, Martinsried, Germany, version 1.5.2.8; http://www.maxquant.org). Different spectrum- and quality-based filter criteria can be then applied to restrict the analyses to highly reliable PSMs only (Tab. 1). In the next step, all potential coding sites (DNA matches) are identified for each high-quality PSM within the given genome sequence considering the degenerated nature of the DNA code. On the basis of the DNA matches found, potential ORFs are deduced according to the following rules: first, an ORF must contain all successive DNA matches encoded on the same strand and in the same reading frame and should not be separated by an interposed stop codon. Secondly, the ORF is extended until the first downstream stop codon encoded in the same reading frame. In the final step, predicting the translational start site, the most upstream DNA match covered by the potential ORF plays an essential role. Three different cases can be distinguished here: (1) The identified peptide encoded by the most upstream DNA match is not of a proteolytic (e.g. tryptic) origin. In this case, the codon directly upstream of the DNA match is assumed to be the translation start site. Since N-terminal methionine residues are cleaved from a number of bacterial proteins during maturation, the translational start is shifted 3 bp upstream if directly preceded by a methionine encoding codon. (2) If the identified peptide encoded by the most upstream DNA match is of proteolytic origin, the assumed ORF is extended to the first upstream codon that is encoded in the same reading frame and directly preceded by a stop codon (primary start). ORF variants then are built starting from each (alternative) start codon between primary start and the most upstream DNA match. The different ORF variants are evaluated based on different measures and features such as presence and location of a ribosomal binding site or product length (Tab. S1). Comprehensive information collected on spectrum, peptide, and DNA level (Tab. S2) is stored in different file formats such as CSV and Genbank. The latter can be used for intuitive result visualization using third party software (Fig. 2C). *Pepper* is open-source and freely available under https://gitlab.com/s.fuchs/pepper.

**Table 1.** Filter and ranking options provided by *Pepper.*

| option | description | this study<sup>1</sup> |
| --- | --- | --- |
| **PSM filter** |  |  |
| --minsample <n> | only PSMs identified in <n> or more samples or replicates will be considered | 2 |
| --perrp <n> | only PSMs with a posterior error smaller than <n> are considered | 0.1 |
| --score <n> | only PSMs with a Andromeda score smaller than <n> are considered | 40 |
| --no_msms | PSMs without MS/MS data are not excluded | - |
| --msms <f> | only high-quality PSMs are considered based on MS/MS ions provided in file <f><sup>2</sup> | + |
| --minintcov <n> | only PSMs with a MS/MS intensity coverage of at least <n> are considered | 0.1 |
| --seqfilter <f> | only PSMs referring to a sequence listed in file <f> are considered | - |
| **Coding site filter** |  |  |
| --uniq_only | multiple coding sites for the same PSM are excluded | - |
| --uniq_only_smart | multiple coding sites for the same PSM are excluded if they do not refer to a gene family<sup>3</sup> | - |

<sup>1</sup> filter <value> used in this study; + indicates that the respective option was used without any value; - indicates that the respective option was not used
<sup>2</sup> high-quality PSMs need to be supported by at least one MS/MS spectra meeting one of the following criteria:
- at least 5 consecutive y- or b-ions - at least 4 y- and at least 4 y-ions - at least 4 b- and at least 4 b-ions - at least 4 b- and at least 4 y-ions
<sup>3</sup> a gene family is defined by 100% sequence identity including sequences of different lengths (gene truncations)

**Table 2.** Identified proteins and peptides using different endoproteases

| protease | unique peptides |  |  | identified proteins |  | unique peptides SP100 |  | identified SP100 |  |
| --- | --- | --- | --- | --- | --- | --- | --- | --- | --- |
|  | absolute number | average length (aa) | alkaline<sup>1</sup> (%) | absolute number | alkaline<sup>1</sup> (%) | absolute number | alkaline<sup>1</sup> (%) | absolute number | alkaline<sup>1</sup> (%) |
| Trypsin | 11886 | 13 | 15.5 | 1696 | 22 | 233 | 19 | 116 | 41 |
| LysC | 3472 | 12 | 17.5 | 1256 | 17 | 106 | 14 | 76 | 37 |
| AspN | 6277 | 13 | 6.5 | 1330 | 16 | 107 | 3 | 56 | 21 |

<sup>1</sup> pI ≥ 8

### 2.4. Small protein identification using different MS-based approaches and databases

To identify small proteins in *S. aureus,* cytoplasmic proteins prepared from cells grown in complex medium to an optical density (OD_540_) of 1 and 7 (both in three replicates) were each subjected to different experimental setups for LC-MS-based protein identification. First, two different techniques for pre-fractionation of proteins/peptides were applied: a gel-based and gel-free approach. For the gel-based approach, proteins were separated by one-dimensional SDS polyacrylamide gel electrophorese (1D SDS PAGE) followed by tryptic in-gel digestion (Fig. 3). The gel-free approach was based on tryptic insolution digestion of proteins followed by an Oasis^®^ HLB solid phase extraction-cartridge purification and strong cationic exchange (SCX)-fractionation (Fig. 3). In both methods, the resulting peptide fractions were subsequently analysed by a nanoAQUITY UPLC System coupled to an LTQ Orbitrap Velos Pro mass spectrometer. In addition, we tested the applicability of a second LC-MS system for the identification of SP100: the Orbitrap Fusion MS coupled to a Dionex Ultimate 3000 UPLC System (Thermo Fisher Scientific Inc., Waltham, Massachusetts, USA). For that purpose, the cytosolic protein extracts were pretreated using the gel-based approach (Fig. 3).

**Figure 3.**
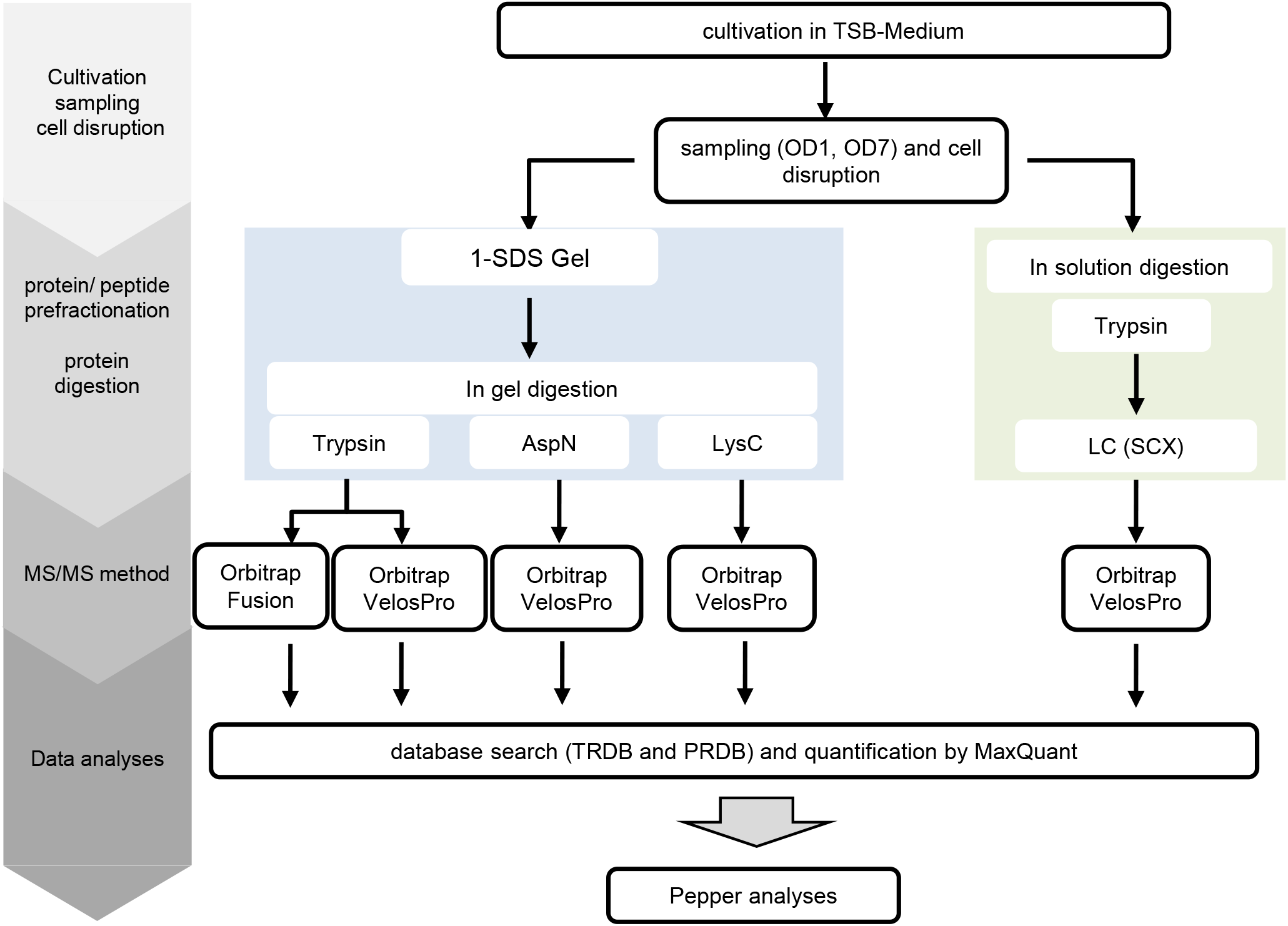
Mass spectrometry (MS) based identification of small proteins in *S. aureus* Newman. The different experimental strategies that have been applied for identification of small proteins (SP100) in *S. aureus* Newman are depicted. Cytoplasmic protein extracts of *S. aureus* Newman grown in TSB medium to an optical density of 1 and 7 were prepared from three biological replicates and aliquots of each replicate were used for the different experimental workflows.

MS- and MS/MS-data of all samples were searched against the new established *S. aureus* databases PRDB and TRDB using Maxquant as an analysis program (Fig. 3). Due to their small size the probability to detect more than one unique peptide for proteins with up to 100 aa is relatively low (see also (21,42). To filter only high-quality unique peptides for protein identification, we modified our MS/MS parameters used by MaxQuant as follows: Andromeda score > 40 for unmodified and modified peptides, delta score > 6 for unmodified peptides and >17 for modified peptides, peptide spectrum matching (PSM) FDR 0.0001. Combining results of all analyses and database searches, 28,153 unique high-quality peptides have been identified, which were further analyzed using our proteogenomics workflow *Pepper* (for spectrum and peptide filtering measures). To minimize false positives, an additional filtering of the MS/MS spectra was applied by *Pepper* (Tab. 1) resulting in 22116 unique peptides that fulfilled the stringent MS/MS quality criteria. These peptides were subsequently mapped to the genome sequence of *S. aureus* Newman. Peptides, which mapped more than once and were not assigned to protein families or to the same protein, were removed from the list manually. This resulted in 21635 unique peptides. 21495 of these peptides were allocated to protein sequences derived from already annotated open reading frames of *S. aureus* Newman. Another 17 peptides supported an extended 5’-end of the respective ORF. The remaining 123 peptides matched to protein sequences that had not been considered in the annotation of the *S. aureus* Newman genome sequence to date (NC_009641.1; 07/06/2013).

To focus on not yet annotated proteins, we relied on ORFs with the highest coding potential based on different features and quality measures (Tab. 1, Tab. S1). In this way, 69 ORFs have been newly described for *S. aureus* Newman ranging between 30 and 346 aa. Focusing on small proteins, we identified 181 proteins of which 65 were not covered by the used genome annotation.

### 2.5. Comparing the applied strategies for small protein identification

To identify which of the applied methods and databases is most powerful for identification of SP100, the obtained identification rates were evaluated. Comparing gel-based and gel-free pre-fractionation methods showed that both methods support the identification of SP100, but with a slightly better identification yield for the gel-free approach (148 versus 135 proteins). The majority of SP100 (n=116) have been recovered with both pre-fractionation methods, while only a small number of proteins were detected by one method only (Fig. 4, Tab. S3).

**Figure 4.**
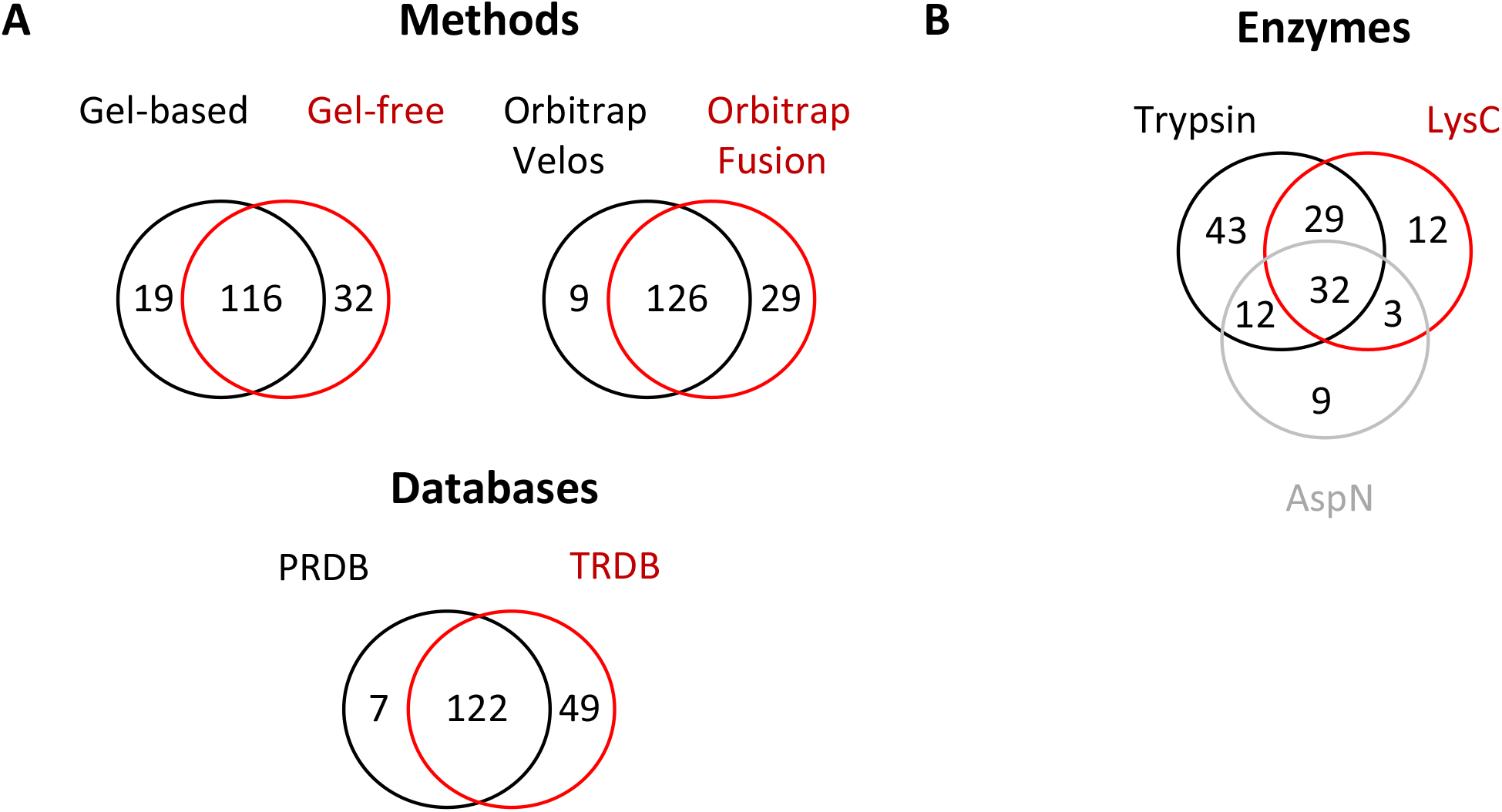
Comparison of the applied experimental strategies for small protein identification. Overlaps of identified SP100 are shown. **(A)** The impact of different prefractionation protocols (gelbased and gelfree), LC-MS instruments (Orbitrap Velos and Orbitrap Fusion MS) and databases (PRDB and TRDB) was analysed. To test the impact of protein/peptide prefraction methods, protein extracts were separated on a one-dimensional SDS gels followed by tryptic in-gel digestion of the proteins within the gel fragments for the gel-based approach. For the gel-free approach tryptic in-solution digestion of proteins followed by an Oasis^®^ HLB-SPE-cartridge purification and SCX-fractionation has been performed. The resulting peptide fractions were subsequently analysed by a nanoAQUITY UPLC System coupled to an LTQ Orbitrap Velos Pro MS. To evaluate the impact of different LC-MS systems, an Orbitrap Fusion MS coupled to a Dionex Ultimate 3000 UPLC System and an LTQ Orbitrap Velos Pro coupled to a nanoAQUITY UPLC System have been used to analyse protein extracts pretreated by the gelbased approach. For MS data analyses, the two *S. aureus* Newman protein databases TRDB and PRDB have been used for each experimental strategy. Finally, considering all experimental strategies, SP100 identified using the TRDB were compared with those using the PRDB. **(B)** To study the impact of different proteases on small protein identification, protein extracts were pretreated by the gelbased approach. In gel digestion has been performed using trypsin, AspN and LysC and the resulting peptides analysed by a nanoAQUITY UPLC System coupled to an LTQ Orbitrap Velos Pro MS. For peptide identification TRDB and PRDB have been used.

The usage of the two LC-MS instruments namely Orbitrap Velos Pro und Orbitrap Fusion in combination with the gel-based approach for protein pre-fractionation led to the identification of 135 and 155 SP100. Out of these, 126 SP100 were identified with both instrument combinations (Fig. 4, Tab. S3). These results suggest that the identification rate of SP100 is higher for the Dionex Ultimate 3000 UPLC - Orbitrap Fusion MS System compared to the LTQ Orbitrap Velos Pro MS based system.

Finally, the impact of both protein databases used (PRDB and TRDB) was evaluated based on the number of identified SP100. With 171 SP100, a significantly higher number of small proteins have been detected using the TRDB, which comprises the entire coding potential of *S. aureus* Newman. In comparison, only 129 SP100 were identified with the PRDB of which only seven were exclusively found with this database. The TRDB was in particular more effective in the identification of proteins not considered in the used genome annotation: while 61 not annotated SP100 have been identified with the TRDB, only 15 were captured using the PRDB (Fig. 4, Tab. S3). This clearly shows, that the ORFfinder prediction algorithm has to be significantly improved. However, out of the 48 not annotated SP100 exclusively found using the TRDB, 29 are encoded by sORFs that overlap with already annotated genes without sharing the same reading frame. Since the prediction classifier has been applied to intergenic regions only, these ORFs could therefore not be found by this approach.

### 2.6 Impact of different proteases on small protein identification

For characterization of the identified SP100, *Pepper* provides several characteristics for each identified protein such as pI and hydrophobicity values (GRAVY). Our data revealed that nearly one third of the identified SP100 (= 58 proteins) depict basic proteins with a pI ≥ 9. This percentage is significantly higher compared to proteins larger than 100 aa where only 17.5% showed these characteristics and is even more impressive when only considering proteins with up to 50 aa (SP50). Here, for almost 50% (n=17) of the identified proteins a pI ≥ 9 was determined (Fig. 5B, Tab. S4). The digestion of proteins by trypsin results in peptides that end with a basic lysine or arginine. In order to exclude any bias, we studied the influence of different endoproteases on the number and pI of the identified SP100. We used the gelbased approach for protein pre-fractionation and the LTQ Orbitrap Velos Pro MS-based system. Besides trypsin, we selected two additional proteases, Lys-C and AspN, for in-gel protein digestion (Fig. 3). MS/MS data analyses have been performed as described above. Using described peptide identification criteria, trypsin digestion resulted in the largest number of peptide identifications (n=11886), followed by AspN (n=6277) and Lys-C (n=3472). The percentage of peptides with a pI ≥ 8 was between 6.5% (for AspN) and 17.5% (for Lys-C (Tab. 2). Next, we examined the number of identified proteins with specific focus on basic proteins (pI ≥ 8). Accordingly, trypsin resulted in 1689 protein identifications of which 22% were basic (pI ? 8). For Lys-C and AspN, we identified 1256 and 1330 proteins and the percentage of identified basic proteins was 17% and 16% (Tab. 2). Accordingly, the impact generated by the use of different endoproteases on the overall identification of alkaline proteins was marginal. However, considering only identified SP100 and relevant peptides, trypsin and Lys-C clearly supported identification of alkaline proteins. With 41 and 37% the percentage is twice as higher than that for all identified proteins. Notably, this is not true for the related peptides. Here the proportion is similar to that found for all identified proteins suggesting that some of the very alkaline SP100 have been identified by acidic or neutral peptides. AspN is the least suited enzyme to identify SP100, although twice as many peptides have been detected compared to Lys-C. This correlates with a significantly lower percentage of alkaline peptides (6.5%) and may support our hypothesis that SP100 tend to be basic. Using Lys-C and AspN, we additionally expected to increase the average length of the identified peptides and therefore enhancing the sequence coverage and the significance of identification results. However, the average length of the identified unique peptides following the digestion with the selected endoproteases was similar, which is in line with observations made by others testing different proteases for proteomics (Tab. 2) (43).

**Figure 5.**
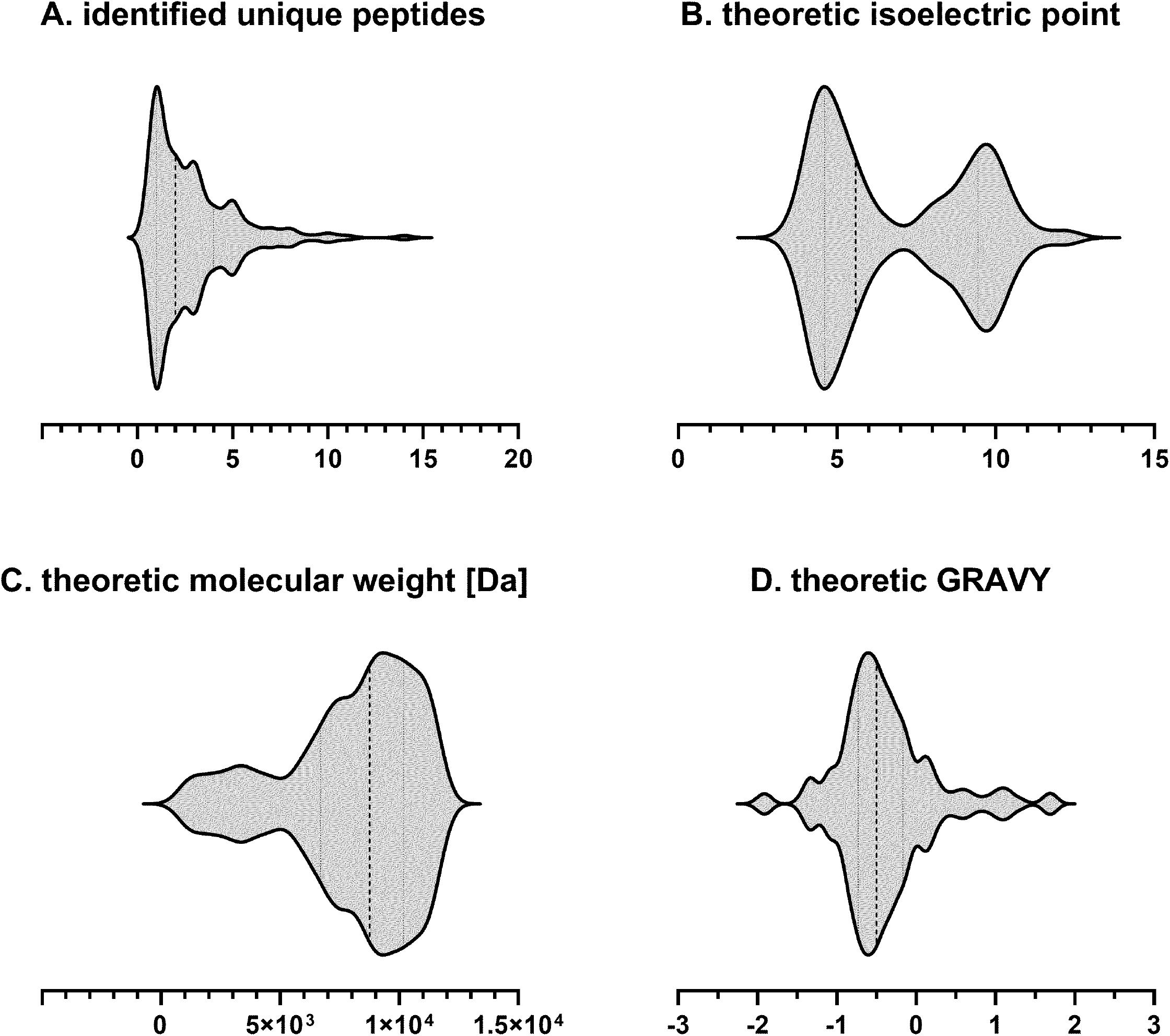
Characteristics of the identified SP100. 185 SP100 have been identified in *S. aureus* Newman using the different approaches. For each protein, the number of identified unique peptides (**A**), the isolelectric point (**B**), the molecular weight **(C)** and the hydrophobicity index Gravy **(D)** was determined. The number of proteins with the respective characteristic shown on the x axis are depicted in violin diagrams.

In total, we identified 141 unique SP100 by using different proteases. As expected, the highest overlap was found for trypsin and Lys-C (n=61 proteins) followed by AspN and trypsin (n=44 proteins). Compared to our methodology approach, only eight SP100 were exclusively identified with the protease approach.

### 2.7. Sequence and expression-based characterization of identified SP100

In total, 185 SP100 were detected using different approaches and the identification of more than half of these proteins (n=110) was based on more than one unique peptide (Fig. 5A). Out of these, 69 were not considered by the used genome annotation of *S. aureus* Newman of which 29 are 50 aa or less. The existence of 154 SP100 (= 83%) was proved by at least two experimental approaches. Among them are 52 newly annotated proteins (Tab. S4). In total, 52% of the identified SP100 (= 96 proteins) had a length of 75 – 100 aa while 28 (15%) possess no more than 50 aa of which 14 proteins are smaller than 26 aa (Fig. 6). To explore possible functions for the uncharacterized proteins we ran BLAST^®^ searches to identify homologs with annotated functions. In addition, we looked for conserved domains using the eggnog database. On the basis of this, we assigned the identified SP100 to functional categories. In total, 142 proteins (78%) were successfully assigned to an eggnog orthologues cluster providing additional support for their existence by biological significance. Out of these, a particularly large set (20%) was linked to translation. This was not very surprising since 16 ribosomal proteins are among the SP100. 16% of the identified proteins were associated with phages, however, their precise role remains exclusive. Another eleven proteins (6%) are involved in virulence of *S. aureus.* These are mainly toxins such as phenol soluble modulins, and delta-hemolysin. Moreover, we found proteins playing a role in transcription, transport and secretion processes. However, there is still a number of proteins (n=31) that have not been assigned to any functional category yet (Fig. 7).

**Figure 6.**
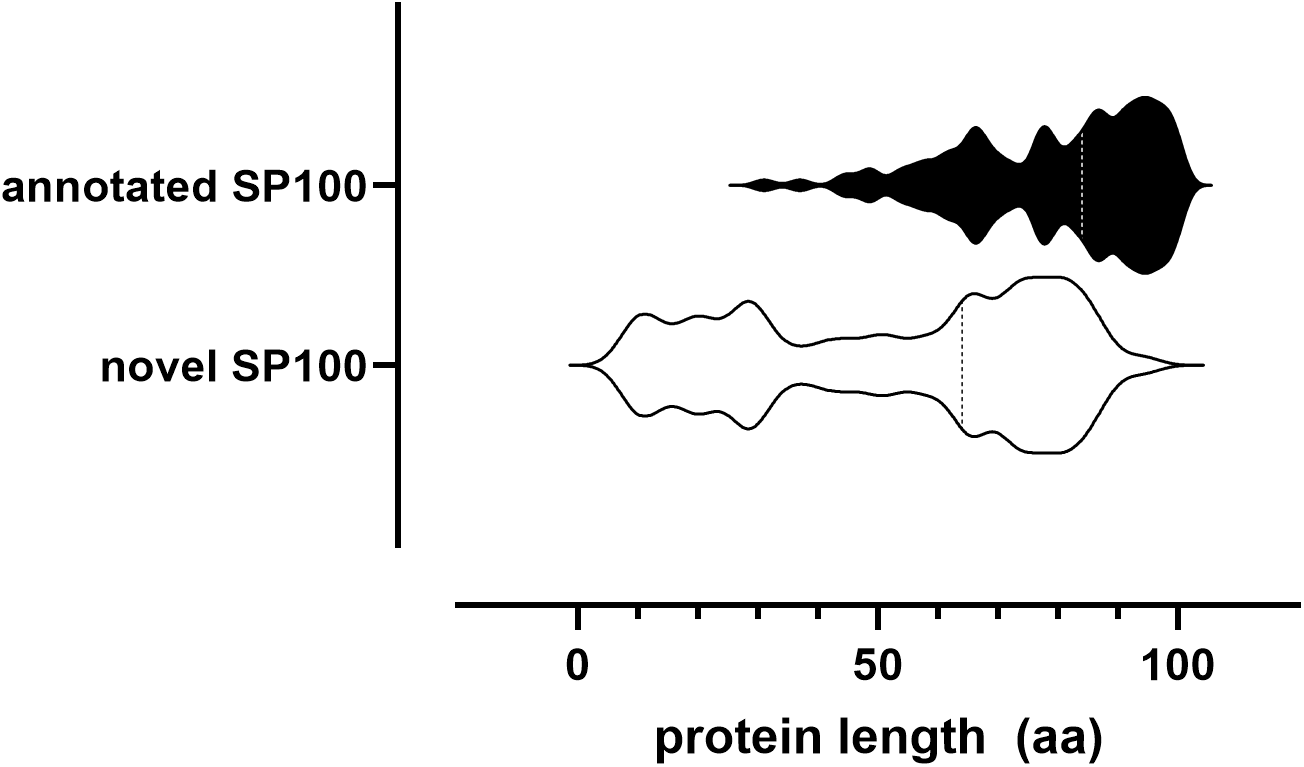
Lengths of identified SP100. The violin plot shows the length distribtuion of annotated and novel SP100 experimentally validated in this study. The respective median is highlighted as dotted line.

**Figure 7.**
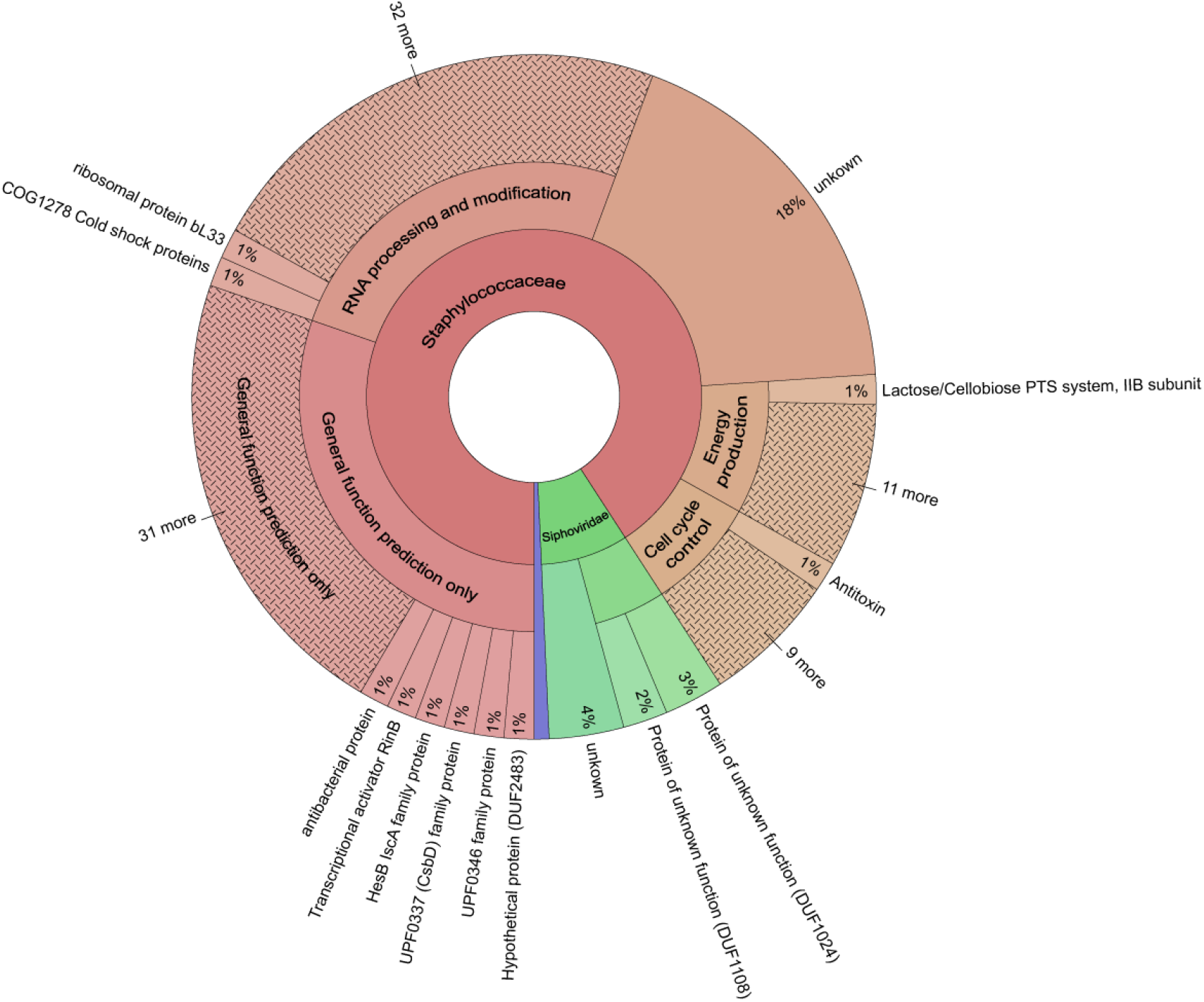
Functional classification of the identified SP100. Conserved protein domains were identified using the eggnog database. On the basis of this, 142 SP100 (78%) were successfully assigned to an eggnog orthologues cluster providing additional support for their existence by biological significance.

To determine whether some of the identified proteins could be secreted or integrated into the membrane we used LocateP (44), which distinguishes seven subcellular localizations of proteins within Grampositive bacteria, and TMHMM Server v.2.0 (http://www.cbs.dtu.dk/services/TMHMM/) providing information on transmembrane domains. This way, in total of 13 of the identified SP100 were predicted be localized outside the cytoplasm. The existence of at least one membrane spanning domain in these proteins strongly suggests that they are integrated into the membrane. For five proteins Sec/SP1 specific-signal sequences were predicted with a probability of 40 to 50% using SignalP 5.0 (http://www.cbs.dtu.dk/services/SignalP/).

A significant drawback of shotgun or bottom-up proteomics is the fact that we can only detect N-terminal peptides when: (i) the translational start site of an ORF is correctly predicted (which is unlikely in the TRDB), (ii) the start codon is preceded by a stop codon or (iii) when it is preceded by a specific cleavage site for the used endoprotease. Only for 17 of the identified proteins, N-terminal peptides were detected. Seven started with methionine while additional three provide evidence that the N-terminal methionine residues were cleaved after translation was initiated. In addition, eight peptides were detected for which the ORF prediction algorithm identified a stop codon immediately in front of the matched DNA region. Notably, although they should therefore represent the N-terminal part of the respective proteins they do not start with methionine. For the remaining sORFs, the true start codon had to be inferred using the prediction algorithm applied by *Pepper.* Accordingly, the majority of the new sORFs has been suggested to start with ATG (n=47). The remaining sORFs presumably initiate at non-canonical translational start sites such as TTG (n=8), ATT (n=3), and GTG (n=2). Interestingly, for eight sORFs, initiation at completely unexpected codons was postulated (Fig. 8). To find further support for the predicted sORFs, we looked for possible upstream ribosomal binding sites. Hence, 48 of the newly derived SP100 encoding genes are preceded by a ribosomal binding site within a distance up to 16 nucleotides upstream of the putative start codon. Notably, out of these 17 shared the sequence (GGAGG with 5 to 9 bp spacer) which is described to be ideal for *Escherichia coli* Shine-Dalgarno sequence (45).

**Figure 8.**
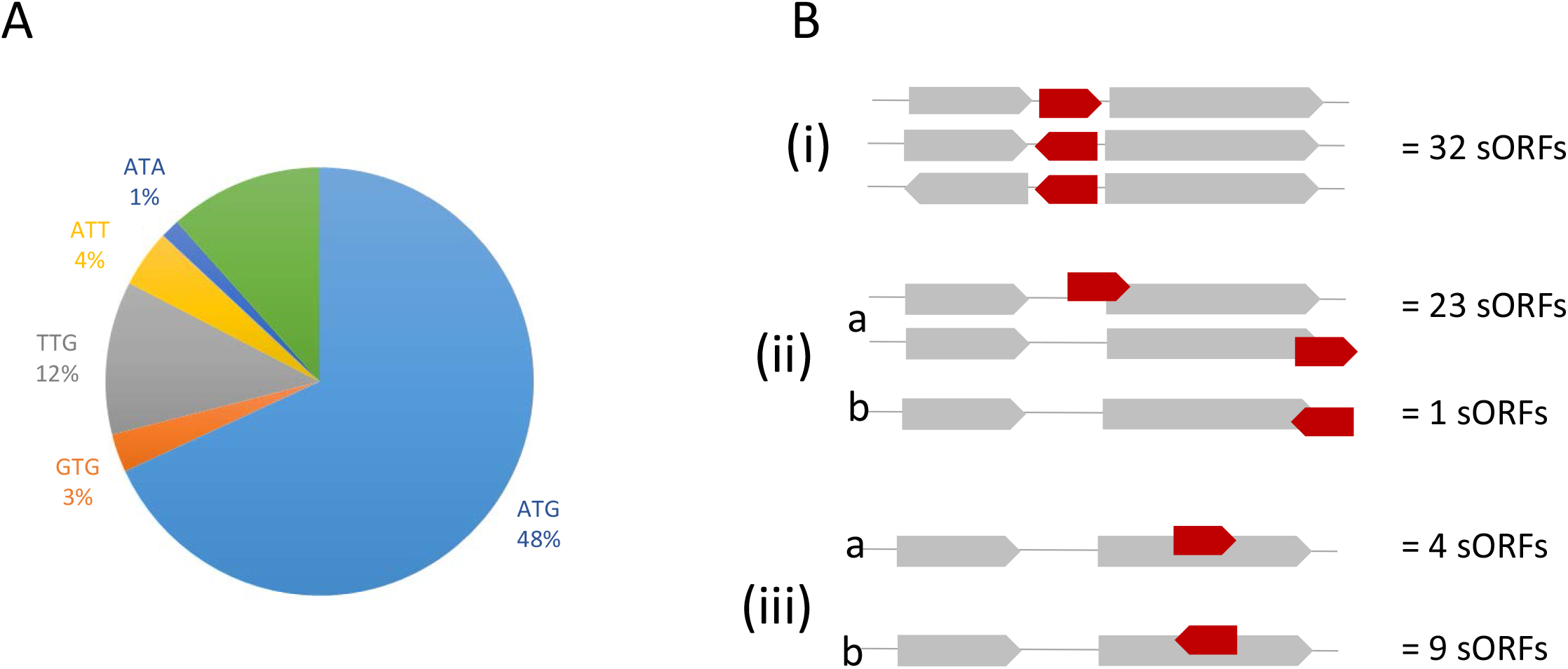
Characteristics of not yet annotated SP100. **(A)** 69 SP100 were identified in *S. aureus* Newman, which were not covered by the used gene annotation. The encoding sORFs were derived by Pepper on the basis of the identified peptides and specific criteria concerning the translational start codon, the Shine Dalgarno sequence and the length of the spacer between both. **(A)** Distribution of different translational start codons between the newly predicted sORFs. **(B**) Characteristics of the genome localization of the newly predicted sORFs: (i) intergenic regions, (ii) partly overlapping with another ORF at the same strand or at the complementary strand, and (iii) completely overlapping with another ORF at the same strand or at the complementary strand.

Furthermore, conservation of these proteins among different *S. aureus* strains as well as among staphylococci was investigated to provide additional evidence for their existence. Accordingly, 40 of the non-annotated proteins are conserved to 95% in at least half of the sequenced *S. aureus* isolates included in this analysis (n=533). Twenty-seven of these proteins are additionally conserved in other staphylococci (Tab. S4; Fig. 9).

**Figure 9.**
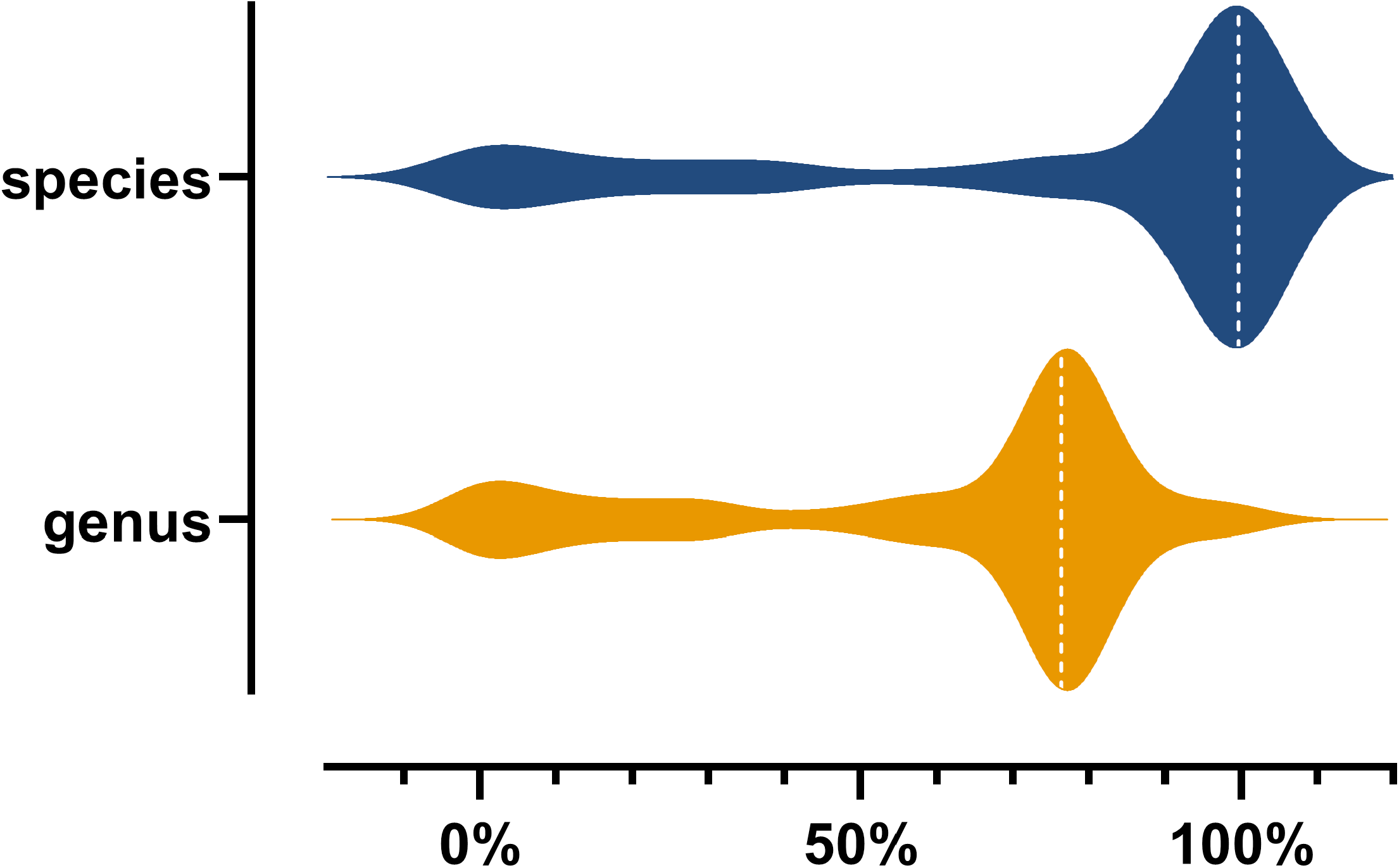
Phylogenetic conservation of the identified SP100 at species and genus level. SP100 were searched against the downloaded genome sequences using tblastn. Based on the best hit alignment to every genome, the identity related to the full query length was calculated. Only alignments sharing 95% identity with the full query sequence were considered. On the basis of these results, relative species and genus conservation rates have been calculated.

Based on their genome localization, the 69 newly discovered sORFs can be classified into three main categories: (i) located in intergenic regions, (ii) overlapping with other ORFs either with the 5’ or with the 3’-end but in a different reading frame, and (iii) located within annotated protein coding sequence but in a different reading frame. For the latter two we can additionally distinguish between those located at the same strand and those located at the complementary strand. As expected, the majority (n=32) belongs to the group (i), but also a considerable number of sORFs was allocated to group (ii) (n=24). However, with four exceptions, the overlap to annotated genes was extremely small and did not exceed eight nucleotides. Notably, 13 SP100 belong to group (iii) of which four are localized at the same strand (Fig. 8).

A label-free quantification approach was applied to identify SP100 in *S. aureus* that are differently expressed during exponential and stationary growth phase in complex medium. Combining MS/MS data from gel-based and from the gel-free approach, 17 SP100 were found to be present at ≥1.5 fold higher amounts at OD 7 compared to OD 1 while 28 SP100 were repressed at higher optical densities. In addition, 13 proteins were detected exclusively at OD 7 and six at OD 1. As expected, SP100 that are significantly down-regulated at higher optical densities do mainly depict ribosomal proteins (n=14) (Tab. S5). Among the induced proteins, we found phenol soluble modulins and SpoVG.

### 2.8. Validation of identified SP100 using ribosomal profiling

To corroborate the existence of the identified SP100 we additionally performed ribosomal profiling for *S. aureus* grown under the exact same conditions. Ribosome profiling provides a snapshot of translation (46) and the position of the translating ribosomes can be assessed with codon precision (47). The sequencing reads, which represent ribosome-protected mRNA fragments (RPF), were mapped to the genome and translation profiles were generated. For 139 sORFs we detected RFPs. Thereby 38 were newly discovered with the presented workflow. For 15 additional sORFs, the obtained results were not significant because of insufficient numbers of reads within the predicted coding region; they would require other validation experiments. In case of sORFs embedded within protein coding regions at the same strand, we were not able to clearly assign RPFs. 29 sORFs, which were identified by the proteogenomics approach, showed no translational activity. Among them are 16 sORFs which are localized on three different lysogenic phages. It is interesting to note, that these phages are highly similar and since we only used uniquely mapped reads (i.e. mapping to only one position of the genome), it is likely that the RPFs were discarded for ambiguity. In sum, we have observed translational activity for the majority (75%) of the sORFs detected by our proteogenomics approach, validating the predictive power of the used MS data sets and translational databases in combination with the here developed proteogenomics tool *Pepper* for data analyses. (Tab. S4).

## 3. Discussion

Prediction and identification of proteins smaller than 100 aa is still very challenging both from computational and experimental point of view. To address this, we here developed a highly optimized and flexible data analysis pipeline for bacterial proteogenomics, covering all steps from (i) protein database generation, (ii) database search, (iii) peptide-to-genome mapping, and (iv) result interpretation. The workflow is based on our bacterial proteogenomics pipeline *Pepper*, extended by *SALT,* a genome translator generating protein and peptide databases using different methods (e.g., stop-to-stop or start-to-stop translation).

To evaluate excellent experimental approaches for identification of SP100, we used a gel-based and a gel-free LC-MS/MS approach in combination with three different endopeptidases. Peptide identifications were obtained by MaxQuant using two different databases: a six-frame “stop-to-stop” translation of the reference genome sequence and the *S. aureus* Newman protein database additionally containing newly predicted proteins using sORF Finder algorithm. Only high-quality peptide identifications were accepted. At the current stage, we prefer the Orbitrap Fusion MS because of its higher sensitivity compared to the Orbitrap Velos MS. Among the prefractionation methods there is no clear favourite. Interestingly, using the 1D gel for protein fractionation, small proteins have been detected in almost all fractions indicating strong interactions with larger proteins. In addition, we tested the value of using different endoproteases for identification of SP100: trypsin, Lys-C and AspN. It became clear that AspN performing hydrolysis of peptide bonds at the amine site of aspartyl residues (48) was less efficient for identification of SP100. Peptides generated by AspN are more frequently acidic. Worthwhile emphasizing is the fact that the pI of 25% of the identified peptides using AspN was below four as compared to only 9% of the identified peptides using Lys-C and trypsin. There is strong reason to believe that the number of basic peptides strongly impact identification of small proteins that tend to be more alkaline. For an improved identification of basic proteins it may thus be essential to consider protocols aiming at higher percentages of basic peptides.

Our combined genome-wide proteogenomics approach enabled us to identify a set of putatively coding sOFRs of which a considerable number has not yet been described. Altogether, 185 soluble proteins with up to 100 aa and 70 of these were not covered by the used gene annotation. 19 of them were identified by at least two peptides. More than 80% were supported by at least two experimental approaches and 139 were actively translated.

The most prominent known small proteins with less than 51 aa in *S. aureus* are phenol soluble modulins with 20 to 40 aa in length (31) and delta-hemolysin with 26 aa (32,49). Interestingly, five of six genes coding for phenol soluble modulins and the delta-hemolysin encoding gene are not present in the used genome annotation of *S. aureus* Newman and thus represent an excellent proof of concept for our proteogenomics pipeline. By applying our stringent identification criteria, we identified three α-type (sORFSaNew0026, sORFSaNew0027, sORFSaNew0028) and two ß-type phenol soluble modulins (sORFSaNew0078, a sORFSaNew0079) and delta hemolysin (sORFSaNew0013) (Fig. S1). With the exception of Psmα3, their cognate sORFs have been correctly predicted by Pepper on the basis of the identified peptides (31,32). For Psmα3 four different ORFs have been derived on the basis of the identified peptides. According to the criteria used for ORF ranking, two sORFs were assigned to orf class 19, one to class 20 and another one to class 21 (Tab. S1). If more than one predicted ORF belongs to the highest-ranked ORF class, *Pepper* favors the larger ORF. Because of valid experimental data regarding the N-terminus of Psmα3 (31), we must unfortunately note that this decision caused a wrongly predicted sORF and insistently shows the limits we still have for the correct annotation of the 5’-end of sORFs. The same might be true for further eight sORFs (Tab.S6). For most of them, however, the predicted ORFs with the same evaluation varied by only a few codons raising the question as to whether multiple start sites may be possible. Additional information provided by ribosomal profiling and MS based identification of N-terminal peptides are thus meaningful to improve ORF prediction. As we have no experimental data for translational start sites, it is certainly the best decision to take the longest ORF. In case of the two sORFs coding for Psmß1 and 2, the predicted N-termini of the proteins were supported by identifying the N-terminal peptides using MS. According to these data, Psmß2 exists in two forms: besides the protein starting with methionine, there is also a processed variant without methionine. We made similar observations for additional 14 small proteins.

Recently, a complete reannotation of the *S. aureus* Newman genome sequence has been performed based on an automated computational analysis using gene prediction method: Protein Homology. As a result, the number of annotated sORFs with less than 100 aa increased significantly. Accordingly, 47 of the sORFs newly identified in the present approach can now be allocated to annotated ORFs in the *S. aureus* Newman data base available from NCBI (NC_ NC_009641.1; 2017-03-02) and the majority overlapped completely with the respective newly annotated genes. However, for 13 newly identified sORFs differences have been observed with respect to the length of the ORFs (Tab. S7). The end of these ORFs is clearly defined by one of the three possible stop codons and with only one exception there are no deviations. In contrast, start sites may vary when multiple start sites are possible and selection of one of these start sites depends on the predicting algorithm used. For selection of the best ORF our analysis tool *Pepper* performs an ORF ranking based on the number of identified peptides, the nature of the start codon, the presence of a ribosomal binding site and the spacer between both. As aforementioned the length of the ORF is only relevant when there are two ORFs belonging to the highest ranked ORF class. Consequently, most of our predicted sORFs are shorter than the corresponding newly annotated ORFs. However, 12 ORFs are preceded by a ribosomal binding site. Of note, two of the sORFs (sORFSaNew0048, sORFSaNew0049) identified by our approach were assigned to a significantly larger gene (NWMN_RS04585), which is a pseudogene with many stop codons within the ORF (Fig. S2). Among the differently annotated genes is also *hld* (sORFSaNew0151). The sORF encoding delta-hemolysin predicted by Pepper results in a 26 aa protein which is in accordance to previously published data (49,50) (Tab. S7).

Notably, one of the identified small proteins with 24 aa in length (sORFSaNew0036) was identical to the N-terminus of the peptide chain release factor encoded by the *prfB* gene in some *S. aureus* isolates. In sequenced *S. aureus* strains the length of the predicted PrfB varies between 291 to 369 aa. In *S. aureus* Newman, the annotated *prfB* gene codes for a 331-aa long protein. Insertion of thymidine at position 810821 results in a frame shift and the ORF starting at position 810751 is prematurely terminated at position 810825, resulting in the mentioned 24 aa containing small protein. Automatic gene annotation identified the ATG codon at position 810866 as an alternative start for *prfB*. Using trypsinolysis, we identified two peptides belonging to the prematurely terminated sORF upstream of the annotated *prfB*. Moreover, we identified one peptide suggesting a 5’extension of the annotated *prfB* gene (Fig. S3) starting at 810803. For both *sORFSaNew0036* and the extended *prfB* gene, translational activity has been confirmed with ribosomal profiling. To exclude that sequence errors may account for the divergent annotation of *prfB* in *S. aureus* Newman, we sequenced the locus and confirmed thymidine insertion at position 810821. Our proteogenomics data suggest translation of two independent ORFs coding for a small protein representing the N-terminal part of PrfB with 24 aa in length and an extended PrfB with 352 aa (Fig. S3). At the current stage we cannot fully exclude that *S. aureus* Newman produces the native PrfB starting translation at position 810751 by reading through the nonsense mutation.

The biochemical roles of most of the identified SP100 are yet to be determined as they still remained largely uncharacterized at molecular level. Colocalization of their encoding genes with already characterized genes, functional domains and localization of the proteins in the cell may provide first hints for the possible role of these proteins in cell’s physiology and virulence. Notably, 16% of the identified SP100 are associated with the prophages. 28 are encoded by φNM1, φNM2 or φNM4, which are members of the siphoviridae family and highly similar. Another two are encoded by φNM3 widely distributed among the human *S. aureus* isolates. Because of the high sequence similarities of φNM1, φNM2 or φNM4, 25 of the identified proteins are orthologues and were associated to 10 protein families. Among them are three orthologous proteins (sORFSaNew0014, sORFSaNew0070, sORFSaNew0134) with similarities to the transcriptional regulator RinB shown to be involved in *int* gene expression (51).

Moreover, we found orthologous proteins containing conserved domains with so far unknown function such as DUF1024 (sORFSaNew0013, sORFSaNew0069, sORFSaNew0135, sORFSaNew0148), DUF1108 (sORFSaNew0066, sORFSaNew0138, sORFSaNew0270) and DUF2483 (sORFSaNew0010, sORFSaNew0137). Functional characterization of these particular proteins is of importance as it has been shown that these prophages play an essential role in *S. aureus* virulence (39) (Tab. S4).

Some sORFs are proximal in sequence space to upstream or downstream genes encoding well characterized larger proteins with enzymatic activity such as peptidase, phosphoglyceroldiesterase, glycolsyl transferase and hydrolase. Hence, it is reasonable to hypothesize that the physiological activity of the newly identified small proteins might be associated with the activity of those larger proteins. Moreover, we identified five SP100 similar to cold shock proteins and four proteins belonging to toxin-antitoxin systems. Most interestingly, almost half of the identified SP100 are basic implying a role in binding to more acidic cellular structures such as nucleic acids or phospholipids. Similar observations have been reported recently for small proteins identified in a simplified human gut microbiome (19). Future work will focus on characterizing these proteins.

Here we present a highly intuitive proteogenomics tool that can be used to identify genes that are hardly predictable by automatic annotation algorithms as for instance very small proteins and to improve gene annotation of already annotated bacterial genomes. In addition, this tool is highly suitable for protein and ORF detection in not annotated bacterial genomes and can support initial genome annotation. Additionally, interesting features such as conservation, pI, GRAVY value, and molecular weight are provided for the resulting proteins (Fig. 5, Fig. 8, Fig. 9, Tab. S4).

## 4. Methods

### 4.1. Bacterial strains, cultivation conditions and cell lysis

*S. aureus* Newman (35) was cultivated in complex medium (TSB) at 37°C and 120 rpm to an optical density at 540 nm (OD_540_) of 1 and 7. Cells were harvested by multiple centrifugation steps and disrupted by cell homogenization (FastPrep-24 ™, MP Biomedicals) (for details see (52)). All experiments have been performed with three biological replicates. The protein concentration was determined using the Roti-Nanoquant assay (Roth, Karlsruhe, Germany) and the protein solution was stored at −20°C.

### 4.2. Fractionation of proteins and peptides and proteolytic cleavage

#### 4.2.1. Gel-based approach

40 μg of cytoplasmic proteins were separated by one dimensional SDS polyacrylamide gel electrophorese (1D SDS PAGE) according to Laemmli (53) with the following modifications: the loading buffer consisted of 3.75% (v/v) glycerol, 1.25% (v/v) ß-mercaptoethanol, 0.6% (w/v) SDS, 0.0014% (w/v) bromophenol blue, 16.5 mM Tris-HCl (pH 6.8). The separation gel contained 12% (w/v) acrylamide gel (with 0.32% bisacrylamide), 0.375 M Tris-HCL (pH 8.8), 0.255% (w/v) SDS, 0.062% (w/v) APS, and 0.062% (v/v) TEMED and the stacking gel 5% (w/v) acrylamide (with 0.13% (w/v) bisacrylamide), 0.125 M Tris-HCl (pH 6.8) 0.25% (w/v) SDS, 0.075% (w/v) APS, and 0.075% (v/v) TEMED.

Proteins were fixed with 40% (v/v) ethanol and 10% (v/v) acetic acid for one hour and subsequently stained with colloidal coomassie (54) for one hour. In-gel digestion using trypsin and extraction of the peptides were carried out as described by Lerch et al. (54) with an additional extraction step using acetonitrile.

For digestion with Lys-C, a buffer containing 25 mM TRIS/HCl and 1 mM EDTA (pH 8.5) was used. The applied enzyme concentration was 1/40 of the total protein concentration. Digestion of AspN was performed in 10 mM Tris-HCl (pH 8.0) with a final AspN concentration of 1/50 of the total protein concentration.

#### 4.2.2. Gel-free approach

The gel-free approach was performed by applying tryptic in-solution digestion followed by an Oasis^®^ HLB-SPE-cartridge purification and SCX-fractionation. In detail, 40 μg of crude protein extract were solved in 8 M urea and 2 M thiourea and adjusted to a final concentration of 6 M urea. After addition of 1.6 μL of 5 mM DTT in 50 mM ammonium bicarbonate solution (pH 7.8), the protein solution was incubated for 30 min at room temperature. For alkylation, 1 μL of a freshly prepared 55 mM IAA in 50 mM ammonium bicarbonate buffer was added to 10 μL of the protein solution and incubated for 20 min in the dark at room temperature. Subsequently, the solution was adjusted to a final concentration of 1 mM CaCl_2_ and 1 M urea using CaCl_2_ solved in a 50 mM ammonium bicarbonate buffer. For digestion, 1 μg trypsin (in 50 mM ammonium bicarbonate and 1 mM CaCl_2_) was applied for 50 μg protein. Digestion was performed for 12 h at 37°C with gentle agitation (50 rpm) and stopped by acidification to a pH value of ≤ 2.5 with 10% formic acid.

For peptide purification, Oasis^®^ HLB-SPE-cartridges (1cc, 10mg, Waters, Milford, MA, USA) were initially conditioned with acetonitrile and then with 0.5% formic acid in 60% acetonitrile. Subsequently, cartridges were equilibrated with two volumes of 0.5% formic acid. Samples were loaded on the cartridges; the flow-through was collected and again loaded on the cartridge. The peptides were washed five times with 0.5% formic acid (FA) and eluted twice with 0.85 ml 60% ACN 0.5% FA. Eluates were dried in a speedvac and frozen at −20 °C.

SCX fractionation was done as described by Kummer et al. (55). To reduce the number of fractions to eight, peptide-containing fractions were combined.

#### 4.2.3. Peptide desalting

ZipTips (C18, Merck Millipore, Billerica, MA, USA) were conditioned with 50% acetonitrile twice. Subsequently they were equilibrated three times with 0.1% FA in 5% acetonitrile. 10 μL of each peptide fraction (resolved in 20 μL 0.1% FA in 5% acetonitrile for 60 min) were loaded on the C18 matrix of the tip by aspirating 10 times. Elution was performed three times by aspirating five times with 0.1% FA in 60% acetonitrile in a new micro test tube. Samples were dried in a speedvac.

### 4.3. Liquide chromatography coupled mass spectrometry (LC-MS/MS)

For LC-MS/MS, each peptide fraction of a sample was solved in 16 μL of 0.1% FA in 3% acetonitrile for one hour, ultrasonicated in a water bath for 5 min and ultracentrifuged.

#### 4.3.1. Orbitrap Velos Pro MS

LC-MS/MS runs with the Orbitrap Velos Pro MS (Thermo Fisher Scientific Inc, Waltham, MA USA) were done as described by Lerch and coworkers (54).

#### 4.3.2. Orbitrap Fusion MS

LC-MS system and used columns are described by Bulitta and coworkers (56). A 200 min gradient was applied, starting with 3.7% buffer B (80% acetonitrile, 5% DMSO and 0.1% formic acid) and 96.3% buffer A (0.1% formic acid, 5% DMSO): 0–5 min 3.7% B; 5–125 min 3.7-–31.3% B; 125–165 min 31.<3–62.5% B; 165–172 min 62.5–90.0% B; 172–177 min 90% B; 177–182 min 90-3.7% B, 182-200 min 3.7% B.

Primary Scans were performed at the Orbitrap in the profile modus scanning an m/z of 350-1800 with a resolution (full width at half maximum at m/z 400) of 120,000 and a lock mass of 445.1200. Using the Xcalibur software, the mass spectrometer was controlled and operated in the “top speed” mode, allowing the automatic selection of as much as posible twice to fourfold-charged peptides in a three-second time window, and the subsequent fragmentation of these peptides. In the non-targeted modus, primary ions (±10 ppm) were selected by the quadrupole (isolation window: 1.6 m/z), fragmented in the ion trap using a data dependent CID mode (top speed mode, 3 seconds) for the most abundant precursor ions with an exclusion time of 13 s and analyzed by the ion trap.

### 4.4. sORF prediction and protein data base generation

We applied nucleotide-composition based sORF prediction and used the sORF Finder algorithm in combination with different training sets. Our positive training set contained all annotated coding sequences from the *S. aureus* subsp. *aureus* strain Newman genome sequence (NC_009641.1; 07/06/2013). In each case the positive training set was combined with sequences from all stable RNAs (sRNAs) encoded by the genome of S. aureus Newman or antisense sequences of all coding sequences as negative trainings sets.

### 4.5. MS Data analysis and statistics

Analyses of the obtained MS and MS/MS data were performed using MaxQuant (Max Planck Institute of Biochemistry, Martinsried, Germany, www.maxquant.org, version 1.5.2.8) and the following parameters: peptide tolerance 5 ppm; a tolerance for fragment ions of 0.6 Da; variable modifications: methionine oxidation and acetylation at protein N-terminus, fixed modification: carbamidomethylation (Cys); a maximum of two missed cleavages and four modifications per peptide was allowed. For the identification of SP100, a minimum of one unique peptide per protein and a fixed false discovery rate (FDR) of 0.0001 for peptides and 0.01 for proteins was applied. The minimum score was set to 40 for unmodified and modified peptides, the minimum delta score was set to 6 for unmodified peptides and to 17 for modified peptides. All samples were searched against the *S. aureus* databases PRDB and TRDB with a decoy mode of reverted sequences and common contaminants supplied by MaxQuant. A protein was considerably reliable identified when it was detected in at least two biological replicates.

For identification of non-annotated open reading frames based on identified peptides a proteogenomics tool has been developed that describes any identified peptide in the degenerated DNA code sequence. By this, exact matches within the reference genome can be found and, subsequently, filtered based on existing annotation and location.

For protein quantification, labelfree quantification mode was selected using MaxQuant LFQ-intensities (57).

Statistical analysis was performed with the Perseus software (Max Planck Institute of Biochemistry, Martinsried, Germany Version 1.5.2.6, www.maxquant.org). LFQ intensities were transformed into log2 values and standardized using Z-score. To evaluate whether the amount of a given protein changed during growth in TSB medium (OD_540_=7 versus OD_540_=1), a permutation-based students t-test (p-value ≤ 0.05) was applied.

A protein was classified to be present exclusively in protein extracts of one of the two growth phases, if it was detected in at least two of three replicates, the raw intensities exceeded the detection limit four times and the raw intensities were below detection limit in all replicates of the respective growth phase.

### 4.6. Phylogenetic and functional analyses

For phylogenetic analyses we downloaded all complete genome sequences for *S. aureus* (n= 533) and staphylococci (n=695) from NCBI RefSeq (state of 03^rd^ March 2020). All SP100 sequences were searched against the downloaded genome sequences using tblastn. Based on the best hit alignment to every genome, the identity related to the full query length was calculated. Only alignments sharing 95% identity with the full query sequence were considered. Based on this, relative species and genus conservation rates have been calculated.

For functional analyses we searched all SP100 sequences against the eggnog database v5.0 using eggnog-mapper 2.0 (default parameter). The taxonomic scope was automatically adjusted to each query to ensure correct classification of phage proteins. Only functions from one-to-one orthology were transferred.

### 4.7 Ribosomal profiling

#### 4.7.1. Library preparation

*S. aureus* cells from 30 ml culture grown in TSB medium to OD_550_=1 were harvested by rapid centrifugation and resuspended in 390 μl ice cold 20 mM Tris lysis buffer pH 8.8, containing 10 mM MgCl_2_ x 6 H_2_O, 100 mM NH_4_Cl, 20 mM Tris (pH 8.0), 0.4 % Triton-X-100, 4 U DNase, 0.4 μl SuperaseIn (Ambion), 1mM chloramphenicol. Cells were disrupted by cell homogenization (FastPrep-24 ™, MP Biomedicals) with 0.5 ml glass beads (diameter 0.1 mm) for 30 s at 6.5 m/s followed by incubation on ice for 5 min. These steps were repeated twice. To remove cell debris, cell lysates were centrifuged and subsequently stored at −80°C and 100 A_260_ units of ribosome-bound mRNA fraction were subjected to nucleolytic digestion with 10 units/μl micrococcal nuclease (Thermofisher) in buffer with pH 9.2 (10 mM Tris pH 11 containing 50 mM NH_4_Cl, 10 mM MgCl_2_, 0.2% triton X-100, 100 μg/ml chloramphenicol and 20 mM CaCl_2_). The rRNA fragments were depleted using the *S. aureus* riboPOOL rRNA oligo set (siTOOLs, Germany) and the library preparation was performed as previously described (58).

#### 4.7.2. Bioinformatic analyses of ribosomal profiling RNAs

Raw sequencing reads were trimmed using FASTX Toolkit (quality threshold: 20) and adapters were cut using cutadapt (minimal overlap of 1 nt) and mapped to the genome version NC_009641.1 (NCBI, January 2020). Following extraction of reads mapping to rRNAs, the remaining reads were uniquely mapped to the reference genome using Bowtie, parameter settings: −l 16 -n 1 -e 50 -m 1 --strata –best y. Non-uniquely mapped reads were non-considered.

## 5. Data Access

The mass spectrometry proteomics data have been deposited to the ProteomeXchange Consortium (http://proteomecentral.proteomexchange.org) via the PRIDE partner repository (59) with the dataset identifier PXD017932.

Ribosome profiling data have been deposited within Gene Expression Omnibus (GEO) under accession number GSE150601.

## 6. Acknowledgements

We thank B. Jung for technical assistance. This work was supported by grants of the Deutsche Forschungsgemeinschaft GRK PROCOMPAS to S.E. and L.J., SPP 2002 (IG 73/16-1) to Z.I., and INST 188/365-1 FUGG to S.E..

## 7. Disclosure Declaration

**Figure S1.**
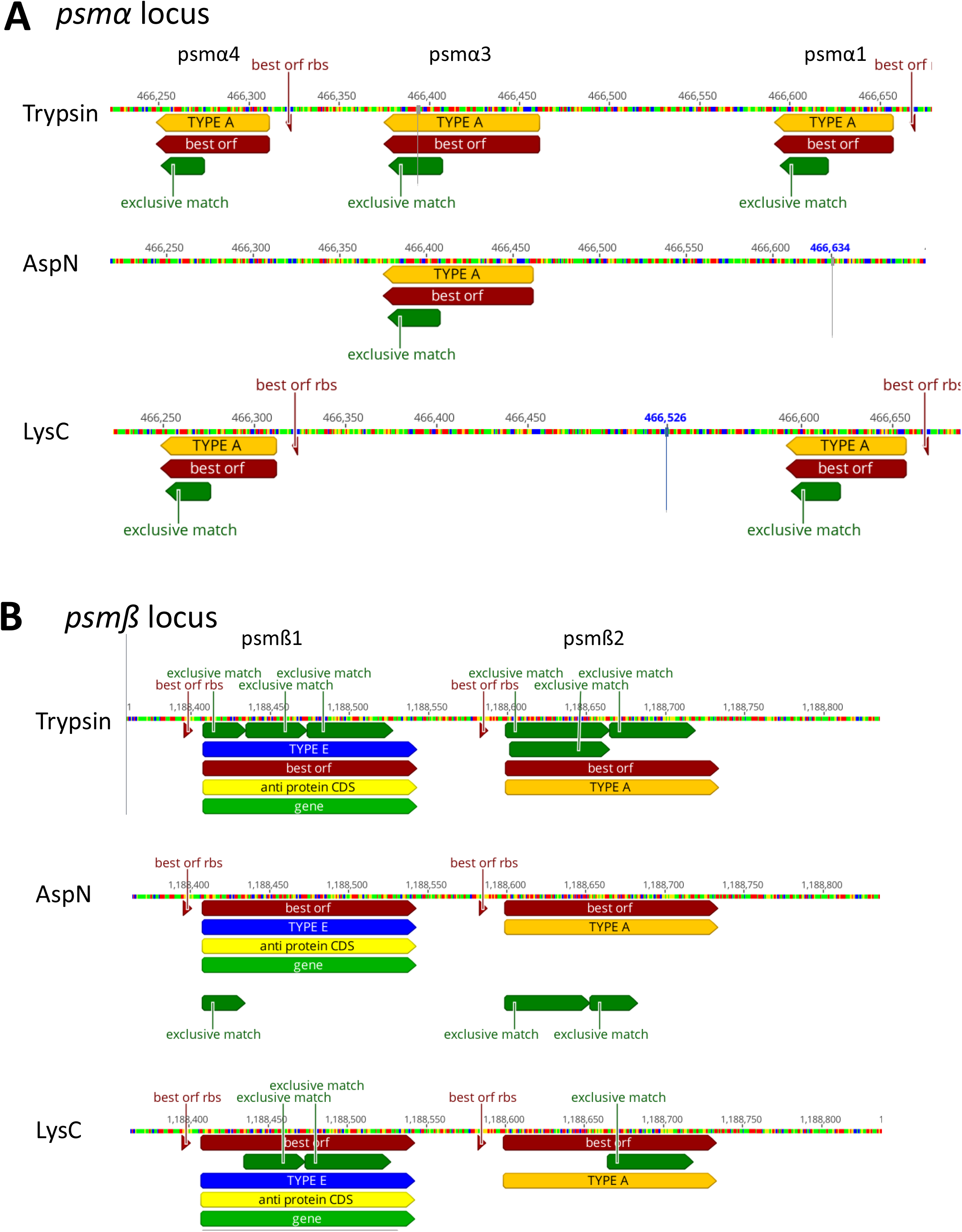
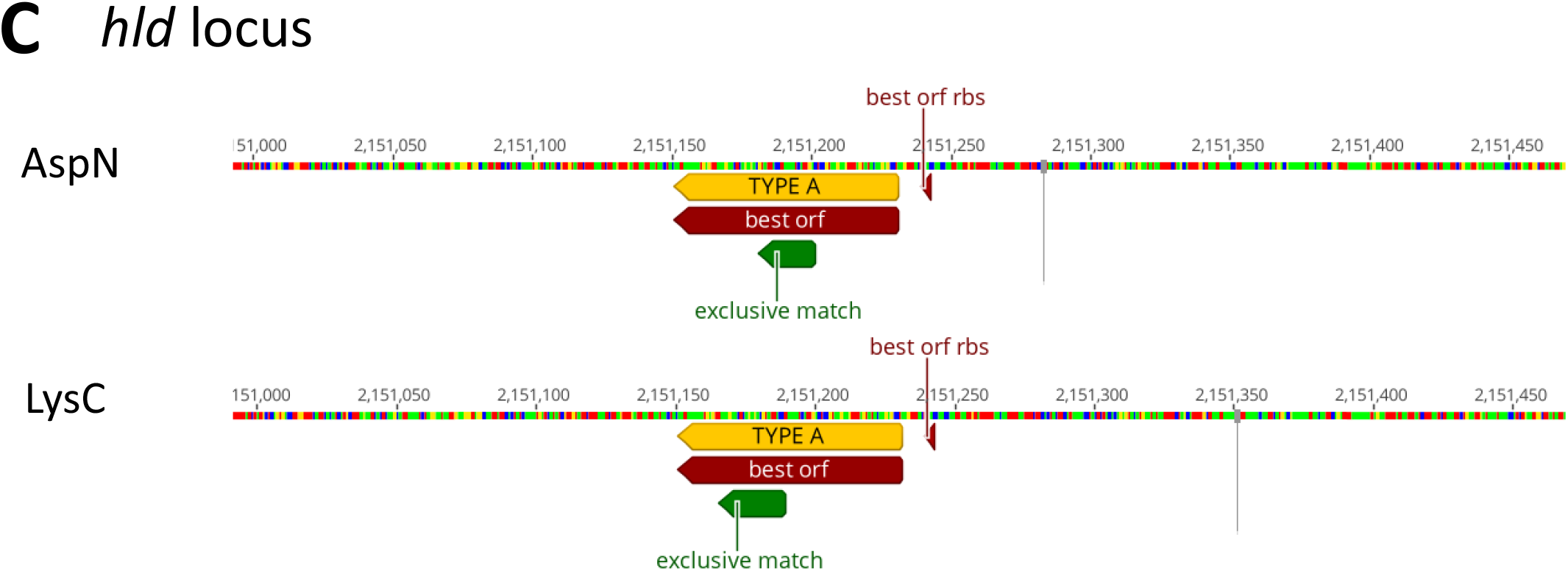
*psma, psmß* and *hld* loci in *S. aureus* Newman. Schematic presentation of the *psma, psmß* and *hld* loci based on the annotation of the *S. aureus Newman* genome sequence (Access: NC_009641.1, BioProject: PRJNA58839, Publication date: 2013-07-06). Annotated genes are shown in light green and the derived coding sequence (CDS) in yellow. Matched unique peptides identified by MS/MS are depicted in dark green and the best ORF derived by Pepper on the basis of the identified unique peptides and additional features is depicted in dark red. In addition, orf types specified by Pepper are presented in orange and blue. Pepper analyses for digestion with AspN, LysC and trypsin are presented separately.

**Figure S2.**
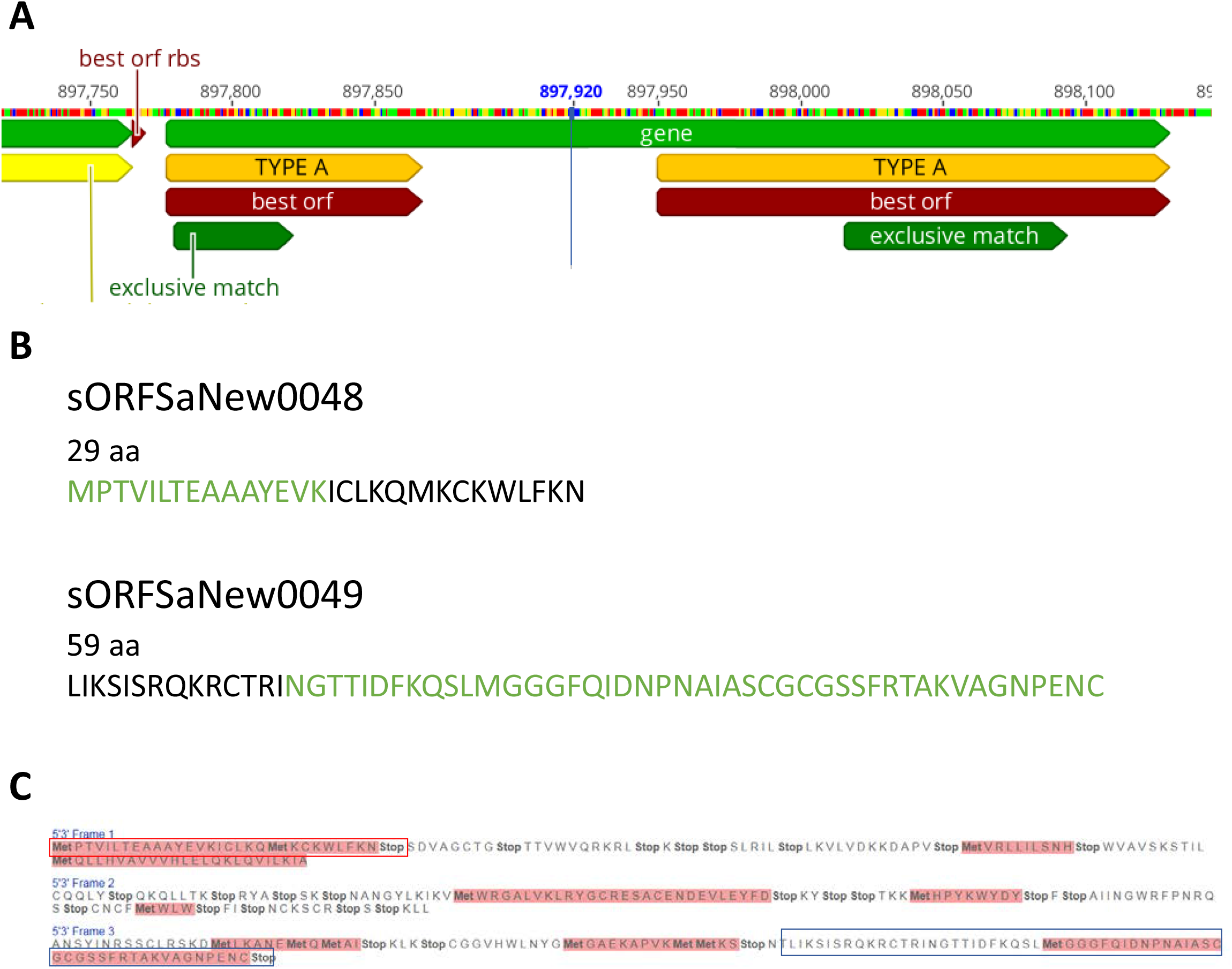
The *NWMN_RS04585* locus in *S. aureus* Newman. **(A)** Schematic presentation of the *NWMN_RS04585* **locus** based on the annotation of the *S. aureus Newman* genome sequence (Access: NC_009641.1, BioProject: PRJNA58839, Publication date: 2017-03-02). Annotated genes are shown in light green and the derived coding sequence (CDS) in yellow. Matched unique peptides identified by MS/MS are depicted in dark green and the best ORF derived by Pepper on the basis of the identified unique peptides and additional features is depicted in dark red. In addition, orf types specified by Pepper are presented in orange. Pepper analyses for digestion with trypsin are presented. (**B**) Protein sequences of the identified proteins sORFSaNew0048 and sORFSaNew0049. Parts of the proteins identical to the original CDS of the pseudogene are marked in green. (**C**) Translational products of *NWMN_RS04585* using all three forward reading frames from stop to stop codon. The coding sequence is interrupted by several stop codons. Translational products corresponding to the identified small proteins sORFSaNew0048 and sORFSaNew0049 are highlighted in red and blue.

**Figure S3.**
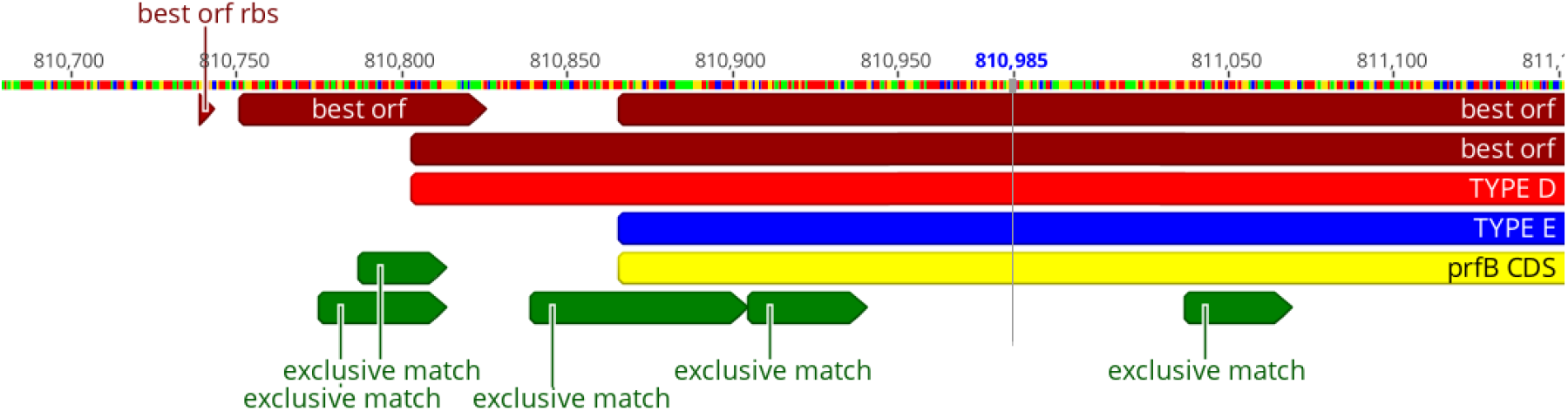
The *prfB* locus of *S. aureus* Newman. Schematic representation of the *prfB* locus based on the annotation of the *S. aureus Newman* genome sequence (Access: NC_009641.1, BioProject: PRJNA58839, Publication date: 2013-07-06). The derived coding sequence (CDS) is shown in yellow. Matched unique peptides identified by MS/MS are depicted in dark green and the best ORF derived by Pepper on the basis of the identified unique peptides and additional features is depicted in dark red. In addition, orf types specified by Pepper are presented in light red and blue. Pepper analyses for digestion with trypsin are presented.

## Notes

### Competing Interest Statement

The authors have declared no competing interest.

